# The crucial role of genome-wide genetic variation in conservation

**DOI:** 10.1101/2021.07.05.451163

**Authors:** Marty Kardos, Ellie Armstrong, Sarah Fitzpatrick, Samantha Hauser, Phil Hedrick, Josh Miller, David A. Tallmon, W. Chris Funk

## Abstract

The unprecedented rate of extinction calls for efficient use of genetics to help conserve biodiversity. Several recent genomic and simulation-based studies have argued that the field of conservation biology has placed too much focus on the conservation of genome-wide genetic variation, and that this approach should be replaced with another that focuses instead on managing the subset of functional genetic variation that is thought to affect fitness. Here, we critically evaluate the feasibility and likely benefits of this approach in conservation. We find that population genetics theory and empirical results show that the conserving genome-wide genetic variation is generally the best approach to prevent inbreeding depression and loss of adaptive potential from driving populations towards extinction. Focusing conservation efforts on presumably functional genetic variation will only be feasible occasionally, often misleading, and counterproductive when prioritized over genome-wide genetic variation. Given the increasing rate of habitat loss and other environmental changes, failure to recognize the detrimental effects of lost genome-wide variation on long-term population viability will only worsen the biodiversity crisis.

## Introduction

Decades of theoretical (1) and empirical (2, 3) research suggest that conserving genome-wide genetic variation improves population viability. Maintaining genetic variation, ***adaptive potential*** (see Glossary) and avoiding ***inbreeding depression*** are central motivations for maintaining large, connected natural populations. Principles of genetics and evolution have therefore played a large role in conservation biology since its inception (4, 5). The genomics revolution has inspired biologists to leverage genome analysis to advance conservation beyond what was possible with traditional genetics. Numerous studies have sequenced genomes of non-model organisms of conservation concern to understand population history, inbreeding depression, and the genetic basis of adaptation. A particularly exciting area of research has been to determine when and how functional genomic information can advance conservation.

Several recent studies suggest that too much emphasis has been placed on genome-wide genetic variation in conservation biology. For example, persistence of small populations for long periods of time despite low genetic variation, and the collapse of the Isle Royale wolf population after the infusion of genetic variation via immigration, have been interpreted as a challenge to the idea that genetic variation generally increases population viability (6–12). Additionally, a weak relationship between conservation status and genetic variation has been used to argue that genome-wide (presumably neutral) genetic variation is of little importance to conservation (11). Several authors have thus advocated for an approach that focuses on functional genetic variation that is thought to directly affect fitness in place of the traditional emphasis on conserving genome-wide genetic variation (6–8, 11).

Here, we evaluate the theoretical and empirical basis of this challenge to the importance of genome-wide variation and show that its premise is inconsistent with population genetic theory and empirical findings. While it is clear that functional genomic information can advance conservation, deemphasizing the maintenance of genome-wide genetic variation would increase the extinction risk of threatened populations.

### 1. Is genetic variation predictive of inbreeding and inbreeding depression?

Inbreeding depression is thought to be driven mainly by homozygous and ***identical-by-descent*** deleterious, partially recessive alleles (13), with lethal and small effect alleles contributing substantially (14). The constant input of new deleterious mutations (15–19) makes ***inbreeding*** depression a ubiquitous phenomenon that can push populations toward extinction (2, 20–23). One of the foundational predictions of theoretical population genetics is that the rate of loss of heterozygosity (***H***) per generation 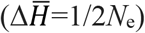 is identical to the rate of increase in mean individual inbreeding (***F***), which is 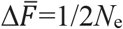 (24). 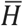 is therefore expected to be entirely predictive of 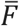 (24–29).

A more difficult, but crucial question is whether genome-wide genetic variation (***π***) is predictive of inbreeding depression. Deleterious alleles are lost in small populations due to selection and genetic drift (30, 31). Deleterious alleles are more often expressed in homozygotes in smaller populations due to inbreeding. Selective ***purging*** of large effect deleterious alleles following inbreeding combined with genetic drift may therefore result in low ***inbreeding load*** and little inbreeding depression in the most highly inbred populations with the lowest π. However, the ability of selection to maintain high fitness in small populations with low π remains unclear.

Population genetics theory predicts that larger populations will have higher neutral (24) and deleterious genetic variation (32, 33). This is illustrated in Figure 1, where simulated large populations have higher *π* (24) and higher inbreeding load (32–34) arising from segregating partially ***recessive*** deleterious alleles (simulations assume empirically supported models of fitness and dominance (***h***) effects; *Supplementary Information* [SI]). Smaller populations have lower π due to genetic drift, and fewer ***lethal equivalents*** due to genetic drift and purging. However, despite having fewer lethal equivalents, chronically smaller populations have lower mean fitness due to partially recessive deleterious alleles being expressed following inbreeding, and some reaching high frequency or fixation (i.e., high ***drift load***). Therefore, a negative relationship is expected between and π drift load for populations at mutation-drift-selection equilibrium.

**Figure 1.**
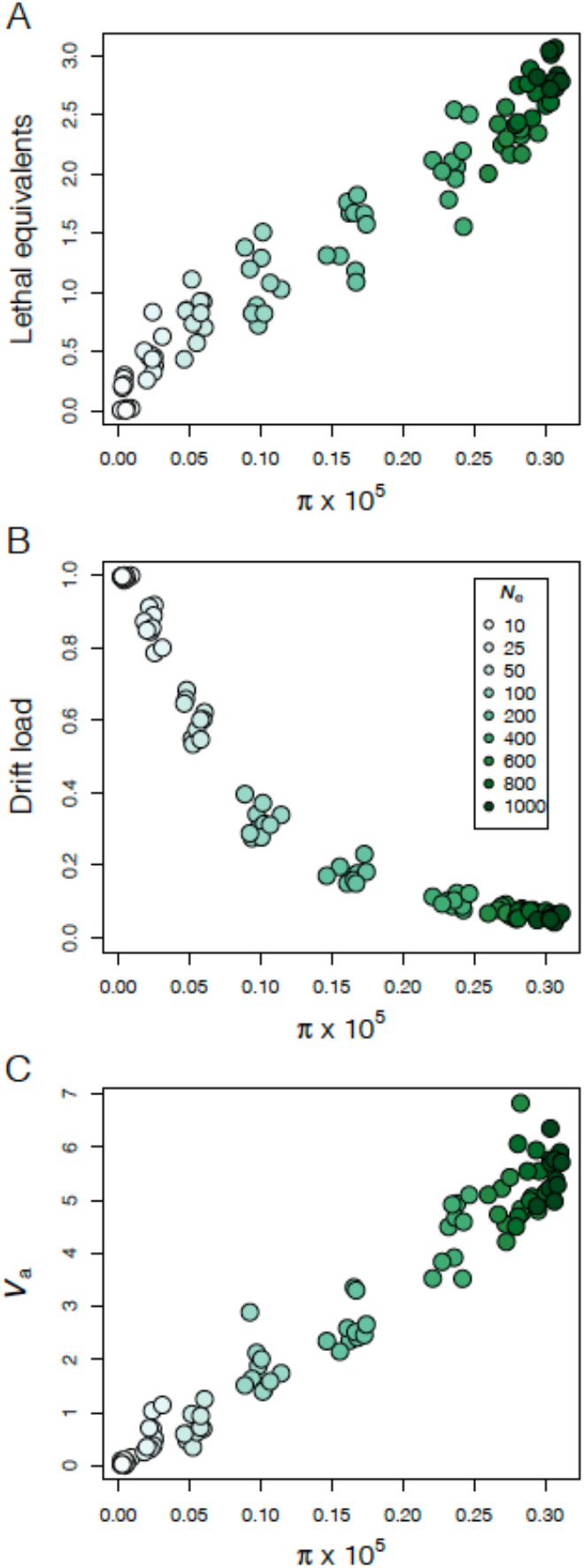
Relationship of nucleotide diversity (*π*) with the inbreeding load (lethal equivalents) (**A**), drift load (**B**), and additive genetic variance in a quantitative trait (*V*_a_) (**C**). The data are from the 1,000^th^ generation of 10 simulated populations with 9 different constant effective population sizes (*N*_e_).

Equilibrium levels of *π* and drift load are not expected in populations with fluctuating population size or immigration rate. A common scenario with high conservation relevance is isolated populations that have experienced recent bottlenecks. The simulated data in Figure 2 shows that genome-wide *π* declines over time following a bottleneck, as expected from classical theory (24) (Figure 2A). This pattern is paralleled by lethal equivalents (Figure 2B) owing to the loss of deleterious alleles via genetic drift and purging of deleterious alleles expressed in homozygotes due to inbreeding (30, 31). However, the deleterious alleles remaining after a bottleneck often go to high frequency or fixation. This results in individuals being homozygous for increasingly more deleterious alleles (higher drift load, Figure 2C) as *π* declines inexorably during a sustained bottleneck, the same pattern expected for small populations at equilibrium (Figure 1). It is notable, though, that *π*, inbreeding load, and drift load can change at substantially different rates following a bottleneck. For example, drift load can become quite high before much *π* is lost following a bottleneck (Figure 2A, 2C). However, small populations that already have low *π* are also expected to have low mean fitness due to ever-increasing drift load, which demonstrates that *π* is a good indicator of drift load and mean fitness. Occasional immigration can be sufficient to maintain high *π* and low drift load in small populations (Figure 2). This is one reason why maintaining connectivity is a priority in conservation biology, and why ***genetic rescue*** is an effective tool for managing small, isolated populations (30, 35, 36).

**Figure 2.**
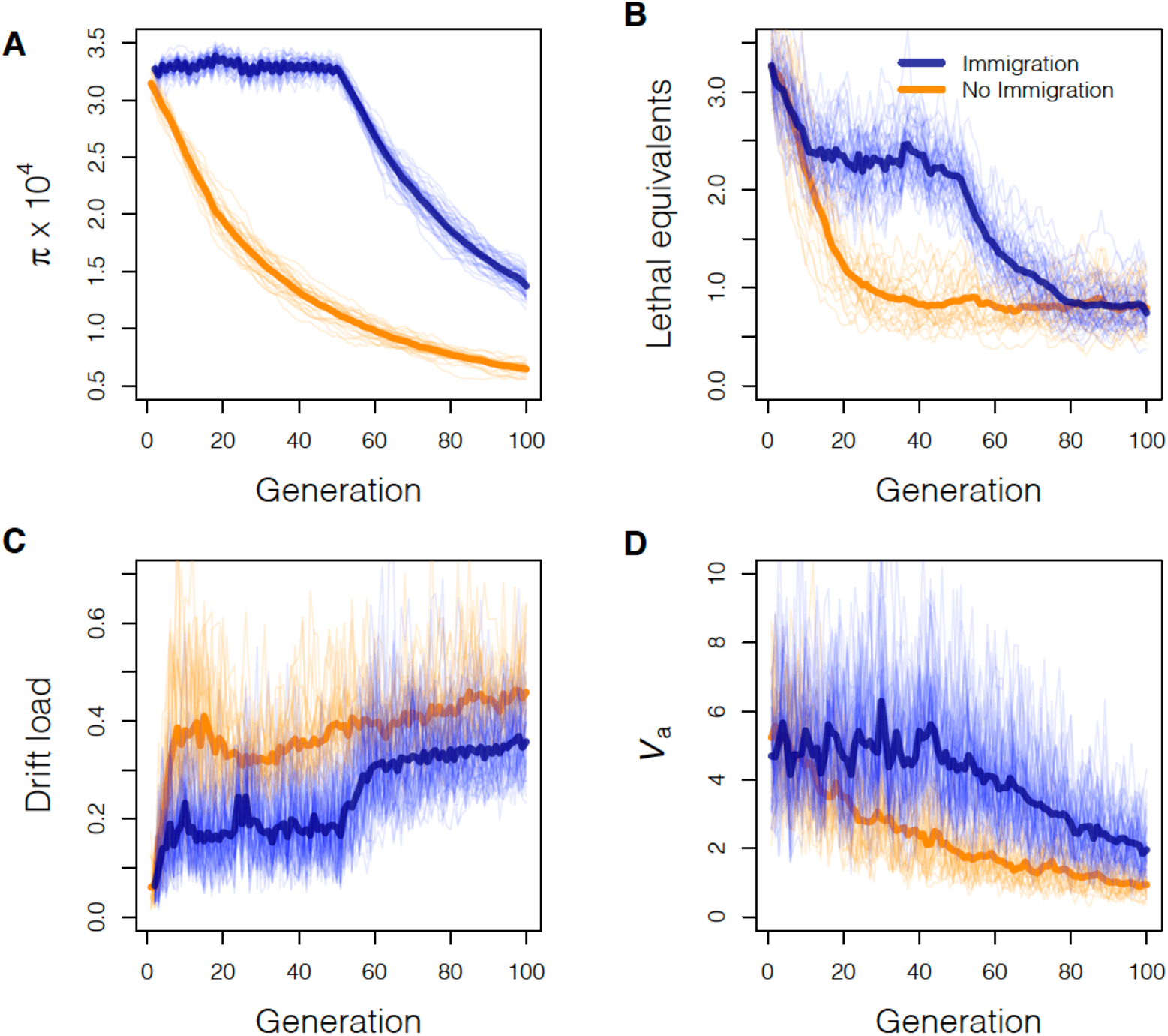
Genetic effects of bottlenecks with and without immigration. Nucleotide diversity (*π*)(**A**), number of lethal equivalents (**B**), drift load (**C**), and the additive genetic variance in a quantitative trait (*V*_a_)(**D**) are shown for 100 generations after a simulated bottleneck in isolated populations (orange) and with 5 immigrants every 2 generations up to generation 50 (blue). Population size was held constant at *N*_e_=1,000 for 1,000 generations before the bottleneck and then at *N*_e_=25 starting at generation 0. The thin lines show the results from 25 replicates. The thick lines represent the mean across 25 replicates. Immigrants during the first 50 generations are from a population with *N*_e_=500 that split from the receiving population the generation of the bottleneck. Details of the simulation model and parameters are provided in the SI.

Empirical data show that purging does not eliminate the extinction threat posed by inbreeding. Pedigree-based studies have yielded mixed results with regard to purging, with typically only a small portion of inbreeding depression being removed after sustained inbreeding in small populations (37–40). Analyses of 60 genomes from seven ibex species found that species which went through the most severe bottlenecks had more deleterious alleles (41). Alpine ibex, which were once reduced to 100 individuals, had fewer highly deleterious alleles but more mildly deleterious alleles compared to Iberian ibex (bottleneck size 1,000 individuals). Empirical genetic data suggest small populations have higher drift load (41–43) which has resulted in lower population growth in populations with lower genetic variation (2, 3). In agreement with theoretical expectations outlined above, these data suggest that purging is insufficient to maintain high fitness in the face of strong genetic drift and inbreeding. Thus, the presence of genomic signatures of purging should not be taken as evidence for the absence of inbreeding depression, or for demographic stability of small populations.

The relationship between *π* and fitness is obviously complicated, particularly immediately after a bottleneck (Figure 2). Populations with the lowest *π* and highest inbreeding will also have the lowest inbreeding load on average due to reduced deleterious genetic variation via genetic drift and purging. However, these same genetically depauperate populations will have lower fitness than larger, genetically diverse populations on average due to ever-increasing drift load (Figures 1 & 2). The bottom line is that reduced fitness is generally expected in small, isolated, genetically depauperate populations due to inbreeding depression and the accumulation of drift load, and that maintaining genetic variation and population connectivity will increase long term viability.

### 2. Is genome-wide genetic variation predictive of adaptive potential?

The ability of populations to adapt to changing environmental conditions (***adaptive potential***) is fundamental for persisting through environmental change (44, 45). A core insight from theoretical genetics is that adaptation requires additive genetic variance (*V*_a_) for the selected trait(s) (46). A lack of *V*_a_ can limit a population’s response to selection and eventually lead to extinction (44, 45, 47, 48). As with other types of genetic variation, *V*_a_ is affected by mutation at loci affecting the trait, selection, migration, and genetic drift (49, 50). We therefore expect from first principles that larger populations will have higher *π* and higher *V*_a_ than small populations *on average* (Figure 1), and thus that *π* should be correlated with *V*_a_. Despite strong theoretical support, determining the strength and importance of this relationship in real populations, especially those of conservation concern, has generated longstanding controversy (51).

Basic population genetic theory shows that population size and connectivity play major roles in determining *V*_a_, and thus adaptive potential. Isolated populations below a certain size should lose *V*_a_ due to genetic drift more rapidly than it is replenished via mutation (49, 52). Additionally, recently bottlenecked populations that have lost *π* will eventually also lose *V*_a_ and evolutionary potential in the absence of immigration (Figure 2). However, while the eventual reduction in *V*_a_ in small populations is inevitable, the initial effects on *V*_a_ following a bottleneck can be complex. Recently bottlenecked populations may show decreases, stability, or even short-term increases in *V*_a_ due to the conversion of dominant or epistatic variance into *V*_a_ as allele frequencies change due to genetic drift (53–55). This potential conversion of nonadditive to additive variation in bottlenecked populations is highly stochastic across traits and populations, and is one of the processes that can cloud the relationship between molecular and quantitative trait variation (56). Nonetheless, the two important takeaways are: 1) although bottlenecks can complicate the prediction of declining *V*_a_ for any given trait in small populations, *V*_a_ will be reduced on average, especially for traits with primarily additive inheritance; and 2) eventually, the inexorable decline in *π* in very small populations means that all small populations will eventually lose *V*_a_ and their ability to adapt to environmental change. Adaptive potential in such populations will be severely limited unless *V*_a_ is replenished by new mutations or migration from differentiated populations (35) (Figure 2).

The hypothesis that small populations harbor less *V*_a_ has been tested empirically in both laboratory and field settings. Most experimental studies show declines in *V*_a_ and weaker responses to selection in small populations or following bottlenecks (57, 58). On the other hand, field studies often find a weak association between *V*_a_ and genome-wide genetic variation when comparing across populations (51, 59); this weak relationship is likely due to a combination of factors, none of which refute the two takeaways described above.

As discussed above, empirical results suggest that *V*_a_ may initially increase after a bottleneck due to the conversion of epistatic and dominance variance to *V*_a_ (54, 60), and then decline after substantial inbreeding accumulates. Further, *V*_a_ is expected to vary among traits and populations depending on genetic architecture, mutation rate, and the mode and history of selection. In practice, most studies are unable to account for these factors and are generally only able to assess a few traits per species/population. Estimates of *V*_a_ for each trait are also typically based on a modest number of families. Although the number of traits, populations, and species studied has increased, determining the total *V*_a_ for fitness in a given population of conservation concern is not an attainable goal. Additionally, the vast majority of the best-characterized species with respect to *V*_a_ in the wild (i.e., most of the species included in (51, 59) meta-analyses) are common. The species and populations in which the relationship between *V*_a_ and genetic variation is expected to be strongest, namely, declining species of conservation concern, tend to be most difficult to characterize.

Arguably the most important point is that the loss of genetic variation in small and/or bottlenecked populations is inevitable and will eventually lead to reduced *V*_a_ and reduce adaptive potential, regardless of short-term and stochastic outcomes. Isolated populations that remain small are unlikely to recover substantial *V*_a_ due to the slow rate of mutation and the counteracting loss of variation to genetic drift, and the lack of adaptive potential is problematic for long term viability (44, 45, 49).

### 3. What is the relationship between genome-wide genetic variation and population viability?

The central question here in relation to conservation is whether populations with lower *π* are less likely to persist. Genetic effects on the persistence of a particular population are difficult to predict with certainty because there are many factors involved that are difficult to evaluate, including mating system and demographic history (32, 33), current and future environmental conditions (61), and the extent to which ***soft selection*** versus ***hard selection*** predominate (62, 63). Additionally, the highly stochastic demography of small populations, which is exacerbated by inbreeding depression (64), means that widely divergent outcomes can be expected across populations with the same environmental, demographic, and genetic starting conditions. However, theoretical empirical studies have yielded broadly applicable insights into the effects of genetic variation and inbreeding on population viability.

Population genetics theory predicts that small, isolated populations with low genetic variation are more likely to go extinct due to genetic effects than larger, more genetically diverse populations under empirically supported mutational assumptions (19, 22, 23, 65). *De novo* mutations following a bottleneck are expected to cause eventual extinction of very small, genetically depauperate populations via ***mutational meltdown*** (Figure S1) (19). The average time to extinction is shorter under the more realistic scenario where bottlenecked populations carry deleterious mutations at the outset (Figure 3). However, the extinction rate depends strongly on bottleneck duration, with longer restrictions conferring increased extinction due to both demographic stochasticity and the constant increase in drift load. Short-lived bottlenecks are one scenario where viability may sometimes be higher for historically smaller, less genetically diverse populations that have fewer deleterious alleles at the outset of the bottleneck due to historical genetic drift and purging (Figures 1, 3A, 3B). However, this assumes inbreeding depression is the only genetic challenge operating, and simultaneous selection caused by environmental change may reverse this relationship. Longer bottlenecks in isolated populations are expected to result in very high extinction rates due to mutational meltdown regardless of the abundance of deleterious alleles at the outset (19) (Figure 3C).

**Figure 3.**
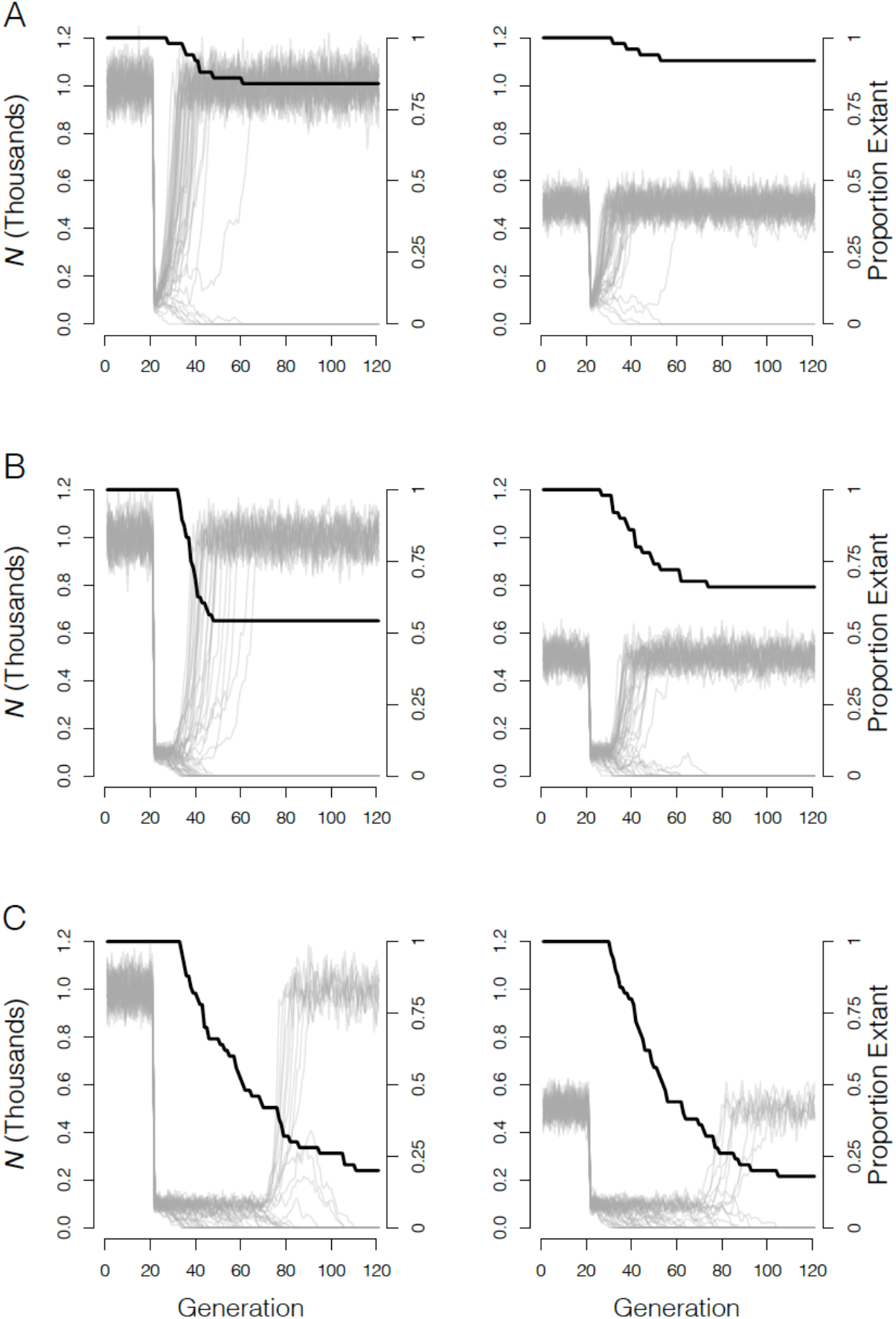
Population viability during bottlenecks from carrying capacity *K*=1,000 (left column) and *K*=500 (right column) to *K*=100 of 2 (**A**), 10 (**B**), and 50 (**C**) generations in length. The black line shows the proportion of extant populations. Gray lines show population size for each of 50 replicate simulations in each scenario.

Empirical studies of population dynamics arguably provide the strongest evidence for the broad benefits of increased genetic variation for population viability. Numerous studies have almost universally found that populations with higher genetic variation have increased population growth and viability. ‘For example’, this relationship has been observed by analyses of the relationship between heterozygosity and population growth rate in alpine ibex (3) and Glanville fritillary butterflies (2). Additionally, the infusion of genetic variation, and the associated masking of previously homozygous deleterious recessive alleles via natural (66) and facilitated immigration (‘genetic rescue’) nearly always increases population growth (35, 36, 67, 68). Although the collapse of the Isle Royale wolf population after a migrant male colonized the population has been interpreted as a counter-example (8), detailed documentation of this population suggested that it was quite extreme and an inappropriate general example because of small island size and the near complete domination of reproduction by one migrant male (67, 69, 70).

Recovery of some populations from severe bottlenecks, and persistence of some populations despite small *N*_e_ and low genetic variation is often cited as a challenge to the idea that low genetic variation and inbreeding reduce population viability (6, 9, 11, 71–74). Soulé (5)[p. 178] pointed out the fundamental flaw of this argument, which he referred to as the “fallacy of the accident” nearly 35 years ago: the only observable populations that have experienced bottlenecks are those that survived. The potentially numerous populations that went extinct are unobservable. Counting extant, genetically depauperate populations is therefore an unreliable metric of the extinction risk posed by lost genetic variation and inbreeding. Theoretical population genetics and population ecology both predict that some populations will survive bottlenecks, and some lucky ones will persist for long periods at small population size. However, such cases are likely the rare exception, the lottery winners so-to-speak, akin to elderly lifelong heavy smokers (5, 75, 76).

The most immediate threats to small, genetically depauperate populations are demographic stochasticity and inbreeding depression. However, long term population persistence will in most cases require populations to adapt to environmental changes (e.g., climate change, novel diseases, invasive species, etc.) (45, 77). Rapid adaptation to new conditions is possible, but requires sufficient genetic variation and relatively large population size (57, 78, 79). All of the material above highlights the fundamental importance of maintaining large, connected, genetically diverse populations. Long term population viability requires having both manageable ***genetic load*** and adaptive potential associated with genome-wide genetic variation.

### 4. What do recent simulation studies tell us about inbreeding depression and genetic rescue?

Simulation results have recently been used to argue that genome-wide genetic variation is of little importance to population viability (8, 11), that inbreeding depression is absent (71), and that genetic rescue is less useful than broadly recognized (8, 11, 71). However, the assumptions in these models are incompatible with our understanding of mutation in real populations, and the best available data on the genetics of inbreeding depression. The deleterious mutation rates per diploid genome (*U*) in these simulations were between 2.6 and 92.3 times lower than in real populations (Table 1). The departure from best estimates from hominids (*U*=1.6) is even greater (16). Further, these simulation studies exclude large-effect mutations (Figure S2). However, highly deleterious mutations occur frequently (15, 16, 80–82), and contribute substantially to inbreeding depression (14, 83–85).

**Table 1.** Deleterious mutation rates used in previous simulation-based analyses of inbreeding depression and genetic rescue.

| Study | Mutation target size | Mutation rate | Proportion deleterious | $U^*$ | $U_{Drosophila}/U^{**}$ |
| --- | --- | --- | --- | --- | --- |
| Teixeira & Huber (11) | 1,000 exons | $1 \times 10^{-5}$/exon/gen | 0.66 | 0.013 | **92.3** |
| Robinson et al. (71) | 2,000 genes × 1,000bp | $1 \times 10^{-8}$/bp/generation | 0.7 | 0.028 | **42.9** |
| Kyriazis et al. (8) | 20,000 genes × 1,5000 bp | $1 \times 10^{-8}$/bp/gen | 0.77 | 0.462 | **2.6** |

\* $U$ is calculated as $2 \times$ mutation target size × mutation rate × proportion of mutations that are deleterious.
\*\* $U_{Drosophila}$ is the deleterious mutation rate per diploid genome, $U=1.2$ , in *Drosophila*(15).

These models clearly produce substantially weaker inbreeding depression than typically observed in real populations. The median number of lethal equivalents for juvenile survival in captive mammal populations is 3.1 (Figure S3) and ranges up to 30.3 (86). Inbreeding load for lifetime reproductive success is rarely measured but expected to be higher than for single traits (87), especially in natural environments (61). The number of lethal equivalents for total fitness under the assumed distribution of fitness effects and models of dominance (***h***) of Kyriazis et al. (8), Robinson et al. (71), and Teixeira and Huber (11) (ranging from <0.05 to approximately 1) are substantially lower than the typical values estimated for a single trait in mammals, and those associated with mutation parameters estimated in model organisms (Figure S3). Thus, these simulations underestimate the fitness effects of drift load and inbreeding depression (71), the importance of genetic variation in conservation (8, 11), the efficacy of genetic rescue as a tool in conservation (8).

### 5. Is the relationship between genetic variation and conservation status informative of the importance of genetic variation to population viability?

It has been suggested that a weak relationship between genetic variation and conservation status (e.g., IUCN Red List) means that genome-wide variation is uninformative of extinction risk (11). However, this relationship is not universally expected, even though extinction risk is strongly affected by genome-wide variation.

First, a lag is expected between reduced population size and the loss of genetic variation. Most threatened populations initially decline due to non-genetic factors (e.g., habitat loss, disease, climate change). Thus, multiple generations are required for a substantial reduction in genetic variation, even after severe bottlenecks (Figure 2A). Threatened populations that became small due to non-genetic factors may still have high genetic variation due to this lag effect. Second, failing to control for other factors that influence genetic variation (e.g., *N*_e_, dispersal, generation time, and mutation rate (11)) is likely to obscure the relationship between genetic variation and conservation status. In contrast, a study controlling for phylogeny (a proxy for the aforementioned confounding factors) has shown a significant relationship between genetic variation and conservation status (88).

Differences among studies in the measures of genetic variation can further obscure true relationships between genetic variation and conservation status. Estimates of genetic variation for different species used in comparative studies vary widely in the number of sampled individuals and populations, and in the regions of the genome analyzed. Some studies estimate species-wide genetic diversity from a single individual (11, 89, 90) and compare different genetic data types across species (6, 90). Using single genomes to estimate species-wide genetic diversity is problematic because the individuals chosen may not be representative of the species as a whole (e.g., captive individuals (89)). Rather, multiple individuals and populations are necessary to accurately reflect a species’ distribution of genetic variation (91, 92). Additionally, estimates of genetic diversity are affected by reference genome quality (93), mapping bias (94, 95), the methods used to measure genetic variation (e.g., whole genome sequencing, RNAseq, RADseq, SNP array, microsatellites), and bioinformatics approaches (92, 93). Thus, sampling, genetic markers, and analyses should be standardized when measuring the relationship between genetic variation and conservation status.

Lastly, IUCN Red List status is an imperfect index of extinction risk. While the IUCN Red List is useful in policy and management decisions, it is a subjective measure of population viability. The IUCN Red List plays an important role in tracking and monitoring the status of biodiversity, but the guidelines and definitions used to categorize threat levels within the Red List are subject to user interpretation, which can lead to inconsistent assessments (96–100). The imperfect relationship between IUCN Red List status and extinction risk means that Red List status is an inappropriate surrogate for extinction risk in assessing the relationship between genome-wide diversity and extinction risk. Together these issues demonstrate that the weak relationship between genetic variation and conservation status has little bearing on the importance of genome-wide variation for extinction risk.

### 6. What is the role of functional genetic variation in conservation?

The widespread availability of genomic data for non-model organisms has rapidly advanced our understanding of the genetic basis and evolution of fitness-related traits in natural populations, e.g., (101–105). This revolution has raised the question of how to effectively integrate functional genomic information into conservation practice (106–109). It has repeatedly been suggested that genetic assessment and management of threatened populations should be focused on variation at particular loci that affect particular fitness traits (11, 110–112). However, such gene-targeted conservation approaches are always difficult, and prone to failure for several reasons.

First, understanding the genetic basis of fitness remains extremely complicated and challenging (106, 108). While some important traits in natural populations are affected by loci with very large effects, most traits are determined by many small-effect loci (113–115). A comprehensive understanding of the genetic basis of such traits is out of reach for non-model organisms (116). To accurately understand the locus-specific effects on a trait and fitness requires information on dominance and pleiotropy, epistasis, genotype-by-environment interactions, and the amount of linkage disequilibrium with other loci influencing the trait or other fitness components (106). These factors are expected to vary among traits and to differ for the same trait among species and potentially among populations within a species, e.g., (101). Therefore, substantial effort is necessary to understand the conservation relevance of a particular genetic variant and predict whether the benefits of gene-targeted conservation actions outweigh potential detrimental effects (106, 108).

A classic example of the potential for undesirable outcomes of gene-targeted conservation management is the suggestion that genetic management of captive and wild populations should be designed around maintaining genetic variation at the major histocompatibility complex (MHC) (11, 110, 111, 117). The MHC has been studied in great detail in humans because of its importance in immunity, organ transplantation, and autoimmune disease, but its organization is unknown or quite different in other vertebrates. Although there is strong evidence for its adaptive importance, some variants have detrimental effects, and the adaptive effects of other variants appear to be environmentally dependent (118). Without detailed examination of the fitness effects of MHC alleles, there is no obvious way to determine fitness effects of particular alleles.

Additionally, as highlighted multiple times over tha last 35 year (119–122)(106, 123) basing conservation management on a small subset of loci risks increasing the loss of genetic variation elsewhere in the genome. Such efforts would be counterproductive unless the gain in mean fitness associated with gene-targeted management is greater than the loss in fitness associated with lost genome-wide variation (106). This highlights the challenges and pitfalls of gene-targeted conservation. When recommendations for maintaining genome-wide genetic variation versus particular adaptive variants are in conflict, a cost-benefit analyses of the two approaches should be performed and a composite solution identified (106). Recent cases where genomic analyses have revealed that large effect loci play a key role in traits of conservation importance, e.g., (101, 102, 104, 124) will be the first to empirically test the efficacy of gene-targeted conservation approaches.

## Discussion

Genomic data should be used to challenge findings from population genetics theory and previous empirical data that form the basis for genetic management of small populations. Recent genomic studies provide useful fodder to determine how to effectively use genomic data to improve conservation in ways that were previously impossible. Examples are emerging of how understanding functional genetic variation could improve recommendations to conserve imperiled populations (101, 102, 104, 124), making genomic data more useful for conservation than ever before. However, genomic data have not discredited the decades worth of evidence that inbreeding depression, mutational meltdown, and loss of adaptive potential are major threats to conservation.

Identifying genetic variants that affect fitness traits undoubtedly advances understanding of the genetic basis of adaptation, and that is important in itself (125). However, placing conservation priority on a small, apparently adaptive portion of the genome ignores what may be the vast majority of variation elsewhere in the genome that will fuel adaptation to unpredictable future conditions (106, 108, 119, 120). This approach is reminiscent of the “adaptationist programme” that Gould & Lewontin (126) criticized >40 years ago for being overly enamored with adaptive explanations for interesting traits (‘spandrels’) without considering that they might have arisen by accident, and that they are but one part of the whole, complex organism (108). Now, as then, we should avoid the temptation to place undue priority on putatively adaptive loci (‘molecular spandrels’ (127)) without first considering the rest of the genome. Our inability to predict future changes in genotype-by-environment interactions should lead us to recognize the importance of genome-wide genetic variation (including presently neutral variation), and more importantly, the factors that make it possible – large livable habitats and natural patterns of connectivity among them.

We know of no convincing evidence that supports abandoning the focus on genome-wide variation in exchange for a focus on functional variation. The recent simulation studies that discount the importance of genome-wide genetic variation in conservation (8, 11, 71) are predicated on genomic assumptions that are inconsistent with the preponderance of empirical data on the genetics of inbreeding depression (see above). Some small populations may not suffer strong inbreeding depression, and some may not rebound following the introduction of genetic variation. However, as pointed out in the formative years of conservation biology, we must resist the temptation to dismiss the extinction risks associated with lost genetic variation in small populations (5).

Although population genetics theory has done a remarkably good job of predicting patterns now observable in genomic data, many questions remain unanswered that will improve the utility of genomic data in conservation. For example, how prevalent is soft selection? The presence of soft selection could help explain some of the instances where populations persist for long periods despite inbreeding (62, 63). How much do *U* and the distribution of fitness effects for deleterious mutations vary among taxa? *U* may be rather consistent within some taxonomic groups (e.g., mammals) where the number of genes is strongly conserved (128). Nevertheless, variation among taxa in gene number, mutation rate, and the amount of intergenic DNA that is subject to deleterious mutation is an important consideration for assessing the fitness effects of inbreeding. Lastly, while it is clear that the distribution of mutation fitness effects is bimodal (80), understanding the specific shape of this distribution, and how much this varies among taxa, is important for our understanding of the extinction risks associated with small population size and inbreeding.

Genomic data will undoubtedly continue to be used to revisit and refine insights gained since genetics was first applied to conservation and to understand the extinction process (4, 5, 47, 129). So far, genomics data have reinforced earlier empirical and theoretical findings showing that genome-wide genetic variation is key to long term population viability. Given the increasing rate of habitat loss and fragmentation, failing to recognize and mitigate the effects of lost genome-wide genetic variation would only exacerbate the biodiversity crisis.

## Acknowledgements

We thank Paul Hohenlohe and Robin Waples for comments on a manuscript draft. DA Tallmon was supported by National Institute of General Medical Sciences of the National Institutes of Health under award number RL5GM118990. WCF was supported by National Science Foundation grants DEB 1413925, DEB 1754821, and DEB 1838282.

## Glossary

*Adaptive potential*: The ability of a population to evolve adaptively in response to selection. Usually measured as narrow sense heritability (the proportion of phenotypic variance attributed to additive genetic effects).
*Drift load*: The reduction in mean fitness of a population due to homozygosity for deleterious alleles.
*F*: The individual inbreeding coefficient: the identical-by-descent fraction of an individual’s genome.
*Genetic load*: The reduction in fitness due to all genetic effects arising from both segregating and fixed deleterious alleles.
*Genetic rescue*: Increase in population growth or reduction in genetic load arising from the immigration of individuals with new alleles.
*h*: the dominance coefficient. A derived allele is recessive when *h*=0 (heterozygous genotypes have the same mean fitness as homozygous wildtypes), and dominant when *h*=1 (heterozygous genotypes have the same mean fitness as homozygous derived allele genotype), and additive when *h*=0.5 (heterozygous genotypes have fitness midway between the alternative homozygous genotypes).
*H*: heterozygous fraction of an individual’s genome.
*Hard selection*: Where an individual’s absolute fitness depends only on its phenotype or genotype and is independent of the phenotypes or genotypes of other individuals in the population.
*Identical-by-descent*: two segments of DNA are identical-by-descent when they both descend from a single haploid genome in recent ancestor.
*Inbreeding*: mating between relatives.
*Inbreeding depression*: reduced fitness of individuals whose parents are related.
*Inbreeding load*: A measure of the potential for inbreeding to reduce fitness, measured by the number of ***Lethal equivalents***, which is a set of alleles that would on average cause one death when homozygous.
*π*: nucleotide diversity: expected proportion of nucleotide differences between randomly chosen pairs of haploid genomes in a population.
*Purging*: Reduction in the inbreeding load owing to deleterious partially recessive alleles being exposed to purifying selection via inbreeding.
*Soft selection*: Selection where an individual’s fitness depends on its phenotype or genotype relative to others in the same population.

## SUPPLEMENTARY INFORMATION

**Figure S1.**
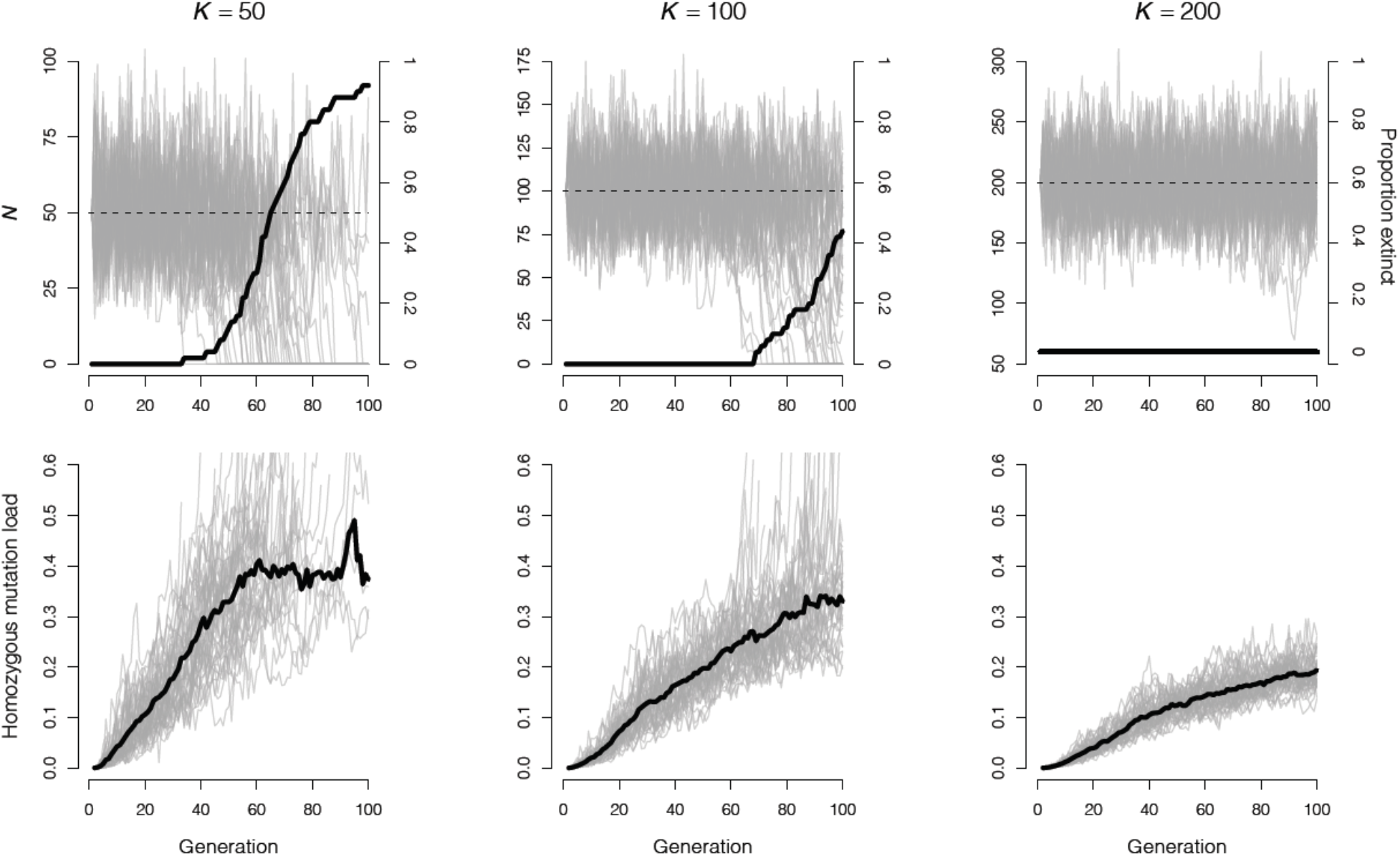
Mutational meltdown via *de novo* mutation in isolated populations. Panels in the top row show population size *N* (gray lines, left vertical axis), and the proportion of extant populations (thick black line, right vertical axis) for 50 replicate simulations of populations with carrying capacities (*K*) of 50 (**A**), 100 (**B**), and 200 (**C**). The bottom row shows the drift load for each simulation replicate (gray lines), and the mean across all non-extinct populations (thick black line). These simulations with hard selection have a ratio of effective population size (*N*_e_) to *N* of approximately 0.25 on average (Figure S5).

**Figure S2.**
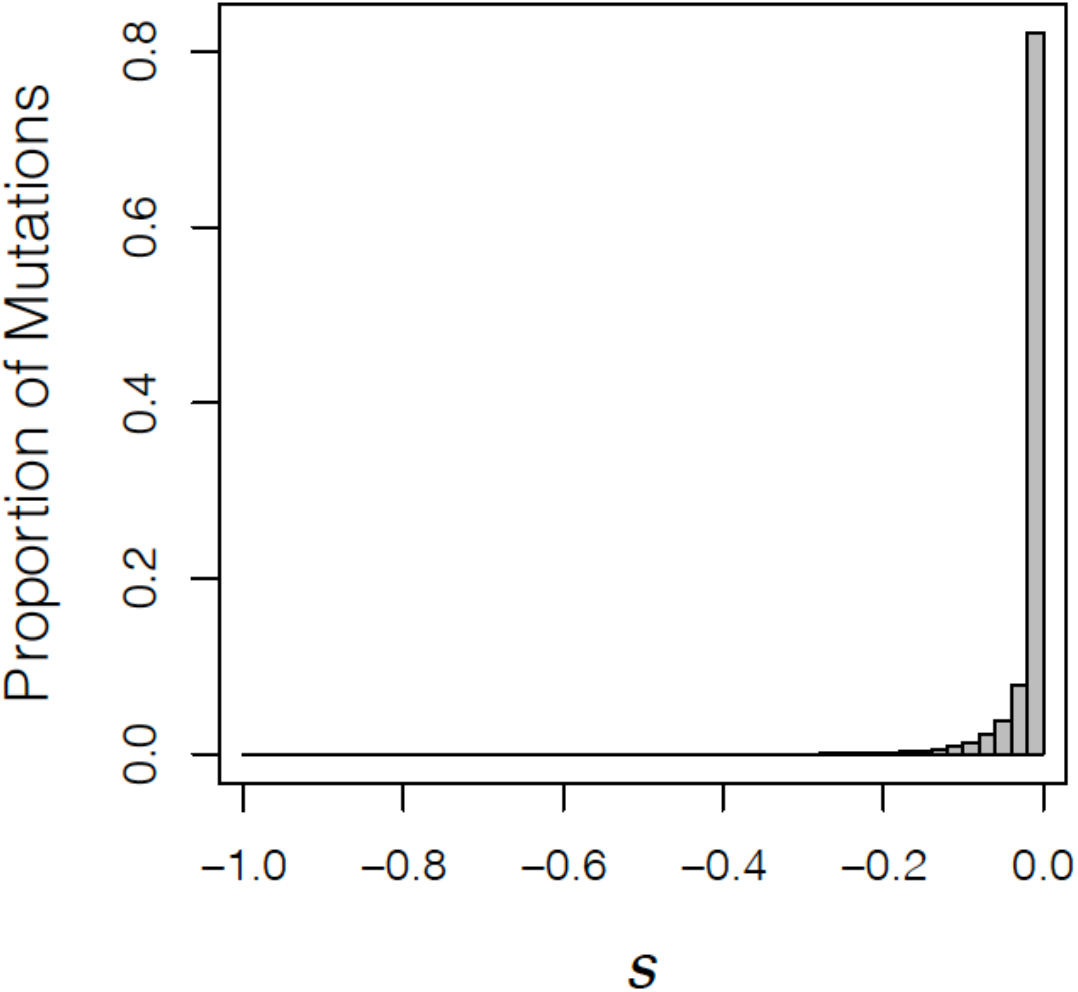
Gamma distribution (shape parameter = 0.186 and scale parameter = 0.071) of fitness effects (*s*) for deleterious mutation assumed in Teixeira and Huber (2021), Robinson et al. 2018, and Kyriazis et al. (2020). Highly deleterious mutations are effectively excluded here (compare to Figure S4).

**Figure S3.**
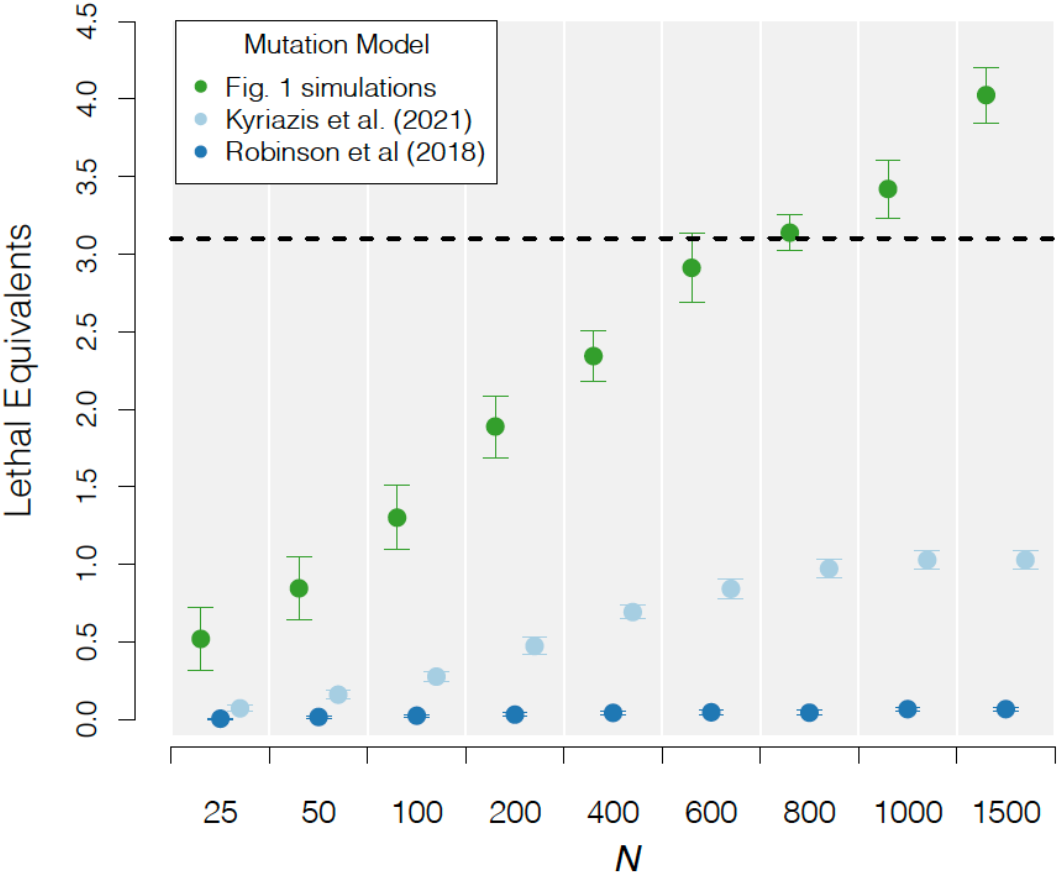
Number of lethal equivalents at approximate mutation-drift-selection equilibrium under the mutation models of Kyriazis et al. (1), Robinson et al. (2), and the simulation model in Figure 1 for constant population sizes ranging from *N*_e_ = 25 to *N*_e_ = 1,500. The error bars represent the standard deviation across 10 simulation replicates.

## Simulations illustrating the relationship between genetic variation and fitness

We use individual-based simulations implemented in R (3–6) to illustrate the relationships among genetic variation, population size, additive genetic variance (*V*_a_), inbreeding load, drift load, and population viability. These are intended to demonstrate patterns that arise directly from population genetics theory under empirically supported combinations of the key parameters. The simulated organism is a self-incompatible hermaphrodite, and has non-overlapping generations, and mean fecundity of 4 (3) when selection was hard (population size is temporally variable), and 2 when selection was soft (population size is temporally constant). Details on the implementation of hard versus soft selection are provided below. Partially recessive deleterious mutations, and mutations that affect the quantitative trait affect fitness by viability selection before breeding when population size is temporally variable (selection is hard), and during the reproduction phase when population size is constant (selection is soft). The simulations in Figures 1 & 2 in the main text include both partially recessive mutations (as described below), and mutations that affect a quantitative trait (also described below). The simulations shown in Figures 3 (main text), S1, and S3 include partially recessive deleterious mutations, but do not incorporate selection on a quantitative trait.

Simulations with temporally variable population size (Figures 3 & S1) assume a ceiling model of density dependent fitness. Here, when population size is > carrying capacity (*K*), mean fitness is penalized so that the expected number of offspring forming the next generation is *K*.

### Mutations affecting a quantitative trait under stabilizing selection

Our model for the inheritance of a quantitative trait is from Kardos & Luikart (3). The quantitative trait is assumed to have an optimal phenotype value of *θ* = 0 (in arbitrary units), a per diploid genome per generation mutation rate of *U*_q_ = 0.147, with phenotypic effects (*a*) drawn from a uniform distribution ranging from −0.5 to 0.5, an environmental variance of *V*_e_ = 4. We assume a Gaussian fitness function:

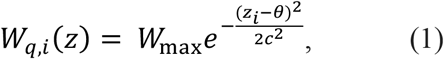

where *W*_*q,i*_(*Z*) is the expected fitness of individual *i* with quantitative trait value *z*_*i*_, *c* is the standard deviation of the fitness function [set to *c* = 6 as in (3)], *z* is the individual’s phenotype value, and *W*_max_ is the expected fitness of an individual with phenotype of *z* = *θ* and no partially recessive deleterious mutations (set to *W*_max_ = 2.5). *W*_max_ is equivalent to the intrinsic population growth rate for a perfectly adapted population with population size very near zero. Smaller values of *W*_max_ (e.g., *W*_max_ = 1.5) resulted in nearly all large populations going to extinction before reaching mutation-drift-equilibrium for lethal equivalents when selection was hard (see below).

### Deleterious mutations affecting fitness

Deleterious mutations act directly on individual fitness. We assume a deleterious mutation rate per diploid genome of *U* = 1.2, as observed in *Drosophila* (7), which is substantially lower than in hominids (*U* = 1.6) (8). The location of a mutation is assigned randomly across 38 chromosomes, the number of autosomes in *Canids* (9), each with a 50 cM genetic length. We assume a gamma distribution of mutation fitness effects (*s*, the expected reduction in fitness for derived allele homozygotes relative to wild type homozygotes), with shape parameter = 0.5 and scale parameter = 0.1, augmented so that 5% of deleterious mutations are lethal (Figure S4). This distribution mimics the distribution of fitness effects observed in mutation accumulation experiments (10), and is consistent with known contribution of both lethal and small-effect, partially recessive mutations in model organisms, humans, and non-model organisms, e.g., (11–13). We assume an exponential model of the relationship between dominance (*h*) and *s* as *h* = 0.5e^−13*s*^, which closely mimics experimental results in model organisms (14, 15), where mutations with *s* very near 0 are generally nearly additive (*h* ≈ 0.5), and mutations with *s* near −1 (lethals) are essentially completely recessive (*h* ≈ 0, Figure S4). Using the higher deleterious mutation rate of hominids would result in an even larger gap between the resulting fitness effects of inbreeding here compared to Teixeira & Huber (16), Robinson et al. (2), and Kyriazis et al. (1) (Figure S3).

**Figure S4.**
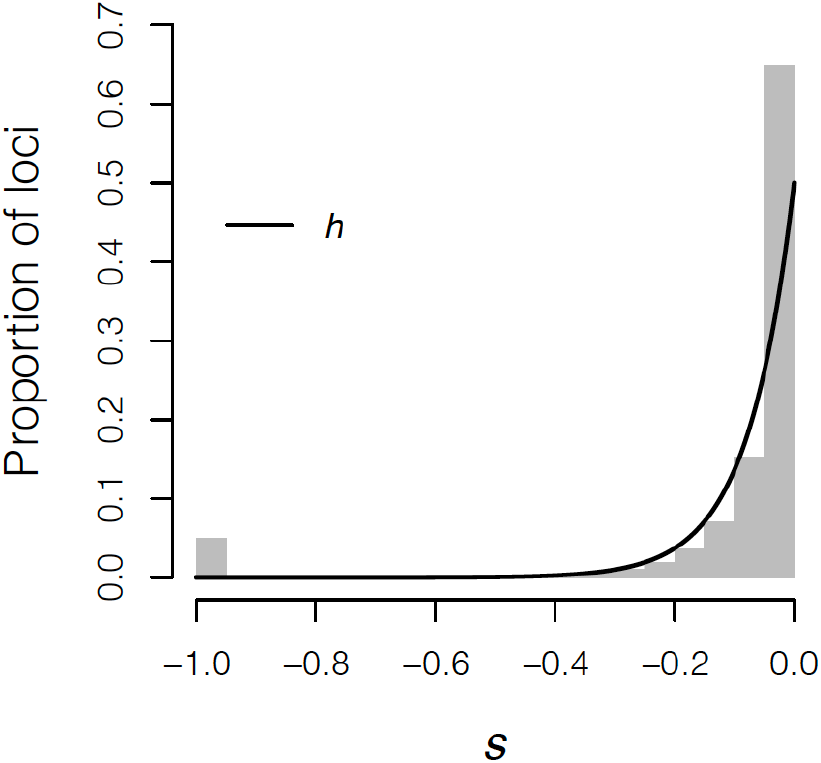
The distribution of selection < coefficients (*s*) for deleterious mutations in our simulations. The black line shows the dominance coefficient *h* as a function of *s*.

The fitness reduction arising from partially recessive deleterious mutations for individual *i* is calculated as

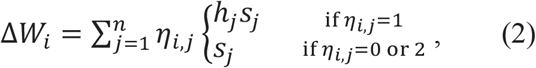

where *η*_i,j_ is the count of the derived deleterious allele at the jth of the *n* loci where there has been a deleterious mutation in individual *i*. *h*_*j*_ and *S*_*j*_ are the dominance and selection coefficients, respectively, at locus *j*. Subtracting Δ*W*_*i*_ from *W*_*q,i*_(*Z*) (eq. 1) yields the expected fitness of individual *i* given the fitness effects of the quantitative trait and partially recessive deleterious mutations.

### Hard versus soft selection

Some of our simulations force population size to be constant (Figures 1, 2, S3) to simplify our analyses of the effects of population size on the parameters of interest. Constant population size implies that selection on the phenotype and arising from deleterious mutations was soft. Here, the mean fecundity is by definition 2, such that the population growth rate is exactly λ = 1, and the expected fitness of an individual with a particular genotype depends on the genotypes of others in the population (17). Selection in these cases is implemented during the reproduction phase.

Our other simulations allowed population size to fluctuate through time (Figures 3 and S1) to illustrate genetic effects on population viability. When population size is allowed to fluctuate through time, selection is hard, where an individual’s fitness depends only on its genotype, and population fitness (population growth rate) depends on the collection of genotypes of all the individuals in the population (17). Here, selection is imposed via selection on juvenile survival before the breeding phase.

**Figure S5.**
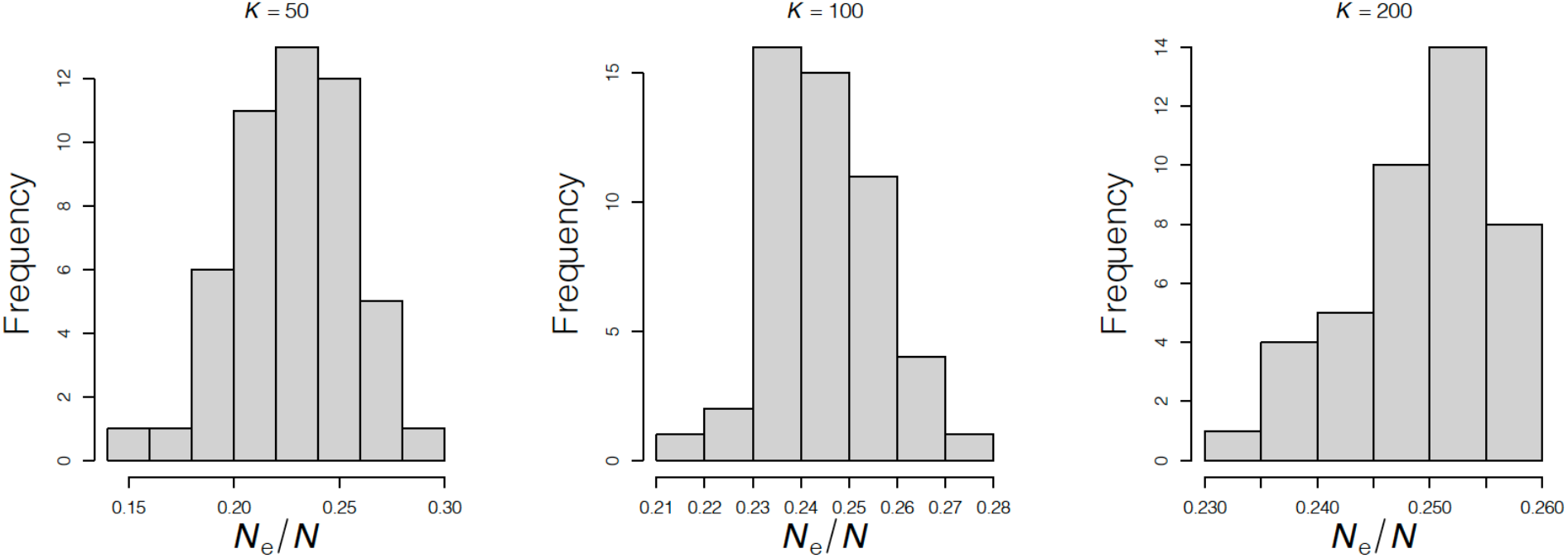
Distributions of the ratio of effective population size (*N*_e_) to census population size (*N*) in simulations from Figure S1. *N*_e_ was calculated as 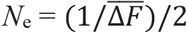, where 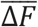 is the mean per generation change in the pedigree inbreeding coefficient in the population over the first <50 generations of the simulation.

